# Impacts of vessel noise on red drum (*Sciaenops ocellatus*) spawning choruses in Saint Andrew Bay, Florida, U.S.A.

**DOI:** 10.64898/2026.02.25.708057

**Authors:** Bennett H. Price, Dakota Brunetti, Amanda Kirkland, T. Erin Cox, Kelly S. Boyle

## Abstract

Noise pollution is an increasing threat to soniferous fishes, however, research on noise pollution impacts is limited to few species and rarely studied *in situ*. Red Drum (*Sciaenops ocellatus*) is an estuarine, fishery species that choruses during spawning. We tested predictions of the hypothesis that Red Drum alter sound production in response to vessel noise. We used passive acoustic monitoring in 2021 and 2022 at an estuarine inlet and Generalized Least Squares (GLS) models to assess vessel sound exposure levels over time (SEL) and other abiotic parameters on Red Drum chorus SELs. GLS models of daily crepuscular choruses indicated a >5% reduction in proportion to crepuscular vessel noise in 2021. GLS models testing influence of abiotic variables and prior vessel noise, predicted reduced chorus SELs proportional to prior noise SEL: ca. 5% and 3% of vessel SEL in 2021 and 2022, respectively. In some instances, SEL during vessel noise was lower than fish chorus SEL immediately prior, indicating instances when fish reduced chorus amplitude during vessel noise or fled the immediate area. In cases when SEL of vessel noise periods exceeded fish calling SEL immediately prior, it is not known if fish modulated calling amplitude because the portion of combined vessel noise and fish chorus amplitude from vessels is unknown. In peak spawning season (September-October) vessel noise was frequent, detected in >31% of recordings in both years and up to 100% of recordings on some dates. Observations of disrupted choruses and high vessel noise prevalence suggest spawning behavior may be impacted by abundant vessel noise.

## Introduction

In aquatic environments, noise pollution affects animal behavior, hearing, and physiology [1–4]. Shifts in behavior include changes in interactions among conspecifics, fight-or-flight response, and modification of acoustic signaling [5–8]. Over recent decades, anthropogenic noise has steadily increased in coastal marine habitats, such as bays and estuaries, with commercial and recreational boat traffic noise representing a major component of this pollution and small watercraft now a ubiquitous source of noise pollution in coastal waters [3,9,10]. Additionally, sound travels farther in water than in air and thus the region of impact of noise pollution is quite large, particularly from sources of low frequency sound [11]. Vessel noise is often low frequency and overlaps with the call spectra and bandwidth of hearing of many fish species [3,12,13], and it is currently believed that all extant fish species can detect low frequency sound [14]. Sound production in fishes occurs in courtship, disturbance, territorial displays and other unknown contexts [15–17]. Disruption of courtship and spawning holds obvious implications for organismal fitness and thus it is important to determine if noise pollution negatively influences these behaviors. Sciaenidae (croakers and drums) present an excellent family for research on impacts of vessel noise on sound production behavior. Sciaenid sound production is associated with spawning and disturbance [16]. Furthermore, members of this family play key ecological roles in estuarine food webs and support economically significant fisheries in the U.S. and globally [18,19,20,21]. Red Drum (*Sciaenops ocellatus*) is a soniferous sciaenid species in which calling is strongly correlated with reproduction [17,22,23]. In the Gulf of Mexico, Red Drum are protected in federal waters and suspected to be overfished, but stock data for Red Drum are limited within this region [24]. Call spectra and hearing range of Red Drum are both low frequency, with dominant frequency of calls between 100-200 Hz and hearing most sensitive from 100-500 Hz [13,25,26,27].

During summer and autumn, Red Drum form large, high-density spawning aggregations associated with intense choruses made by male fish in the afternoon into the evening and peak from August through October [17,23]. Notably, vessel noise can occur in the same habitat and overlap temporally and in the sound spectrum with Red Drum calls [28]. This overlap in vessel noise and call spectra presents a potential risk for acoustic communication with implications for spawning success if fish reduce or terminate calling during vessel noise or if fish continue to call when their signals are unable to be received by conspecifics.

Our study tests for influence of noise pollution on Red Drum sound production behavior in the northern Gulf of Mexico at Saint Andrew Bay in Panama City Beach, Florida, U.S.A. (Fig. 1). This field site was chosen because both mature Red Drum and frequent vessel traffic were observed in this bay by the authors. Therefore, the field site selected afforded an opportunity to test whether vessel noise impacts vocal behavior during the 2021 and 2022 spawning seasons (July-October). Additionally, Saint Andrew Bay is located in the greater Panhandle region of western Florida, where both Red Drum stocks and fishing pressure have been increasing, with fishing effort doubling in this area over the past decade and 2023 having the highest abundance of YOY Red Drum in over 20 years [29].

**Fig 1.**
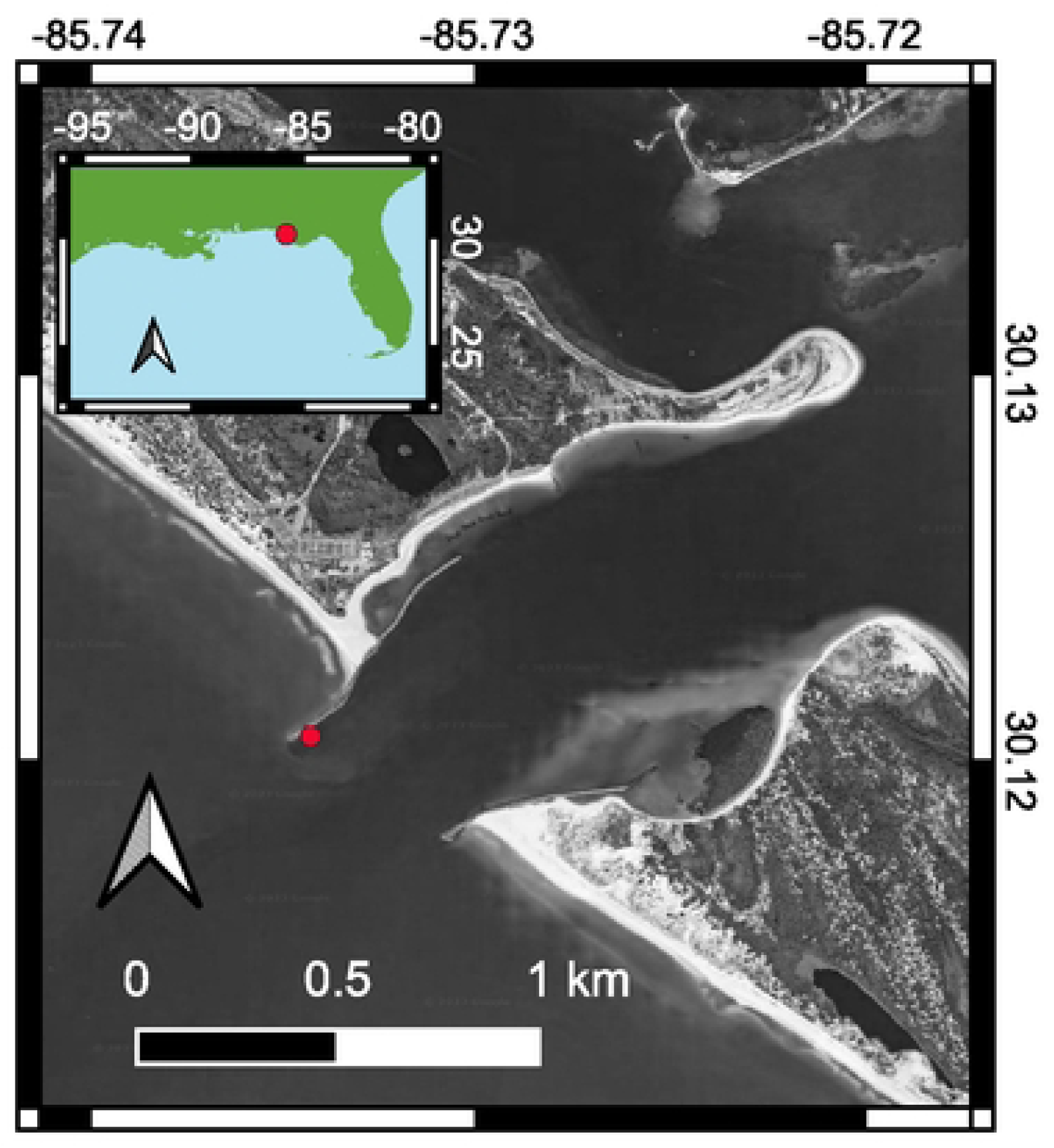
Saint Andrew Bay, Florida, U.S.A. The red star notes placement of the hydrophone on the southwestern end of the jetty at the mouth of the bay (Google satellite imagery, 2025).

We hypothesize Red Drum alter their calling when vessel noise co-occurs in St. Andrew Bay. We considered two alternative outcomes for how soniferous fish may respond in the presence of vessel noise. First, the disturbance hypothesis predicts that fish calling behavior is disturbed by vessel noise. Predictions of this hypothesis are that fish reduce or completely cease calling during and shortly after vessel noise exposure. Consistent with this prediction, in previous studies, a reduction in calling behavior was observed in Sciaenidae and Gobiidae in association with vessel noise [7,8,40]. Additionally, overall spawning success was lower in gobies exposed to noise pollution [7], and in Red Drum, spawning was only observed with increased drumming behavior [17,22]. Fish can also flee the noisy area which would also reduce drumming behavior and be consistent with the disturbance hypothesis. As such, reduction in sound production due to a disturbance response may adversely impact reproductive success.

Alternatively, we considered a compensation hypothesis that predicts fish will increase or alter calls to compete with higher levels of background noise. This behavioral shift of call amplitude or dominant frequency in the presence of excessive background noise is known as The Lombard Effect [30,39]. Of note, even if an animal alters its signal, employing The Lombard Effect, to be detectable through background noise, calls may still be masked when background noise is high, which would result in wasted signaling effort.

The null hypothesis in our study would include no-modification of fish sound production in the presence of vessel noise. This would mean fish call at the same intensity during vessel noise as they do in periods without. Of note, even if fish do not moderate calling behavior, if calls are masked and not detected by the intended receiver in a spawning aggregation, calling could result in wasted effort and there may be adverse impacts on reproduction.

Our study examined Red Drum calls and vessel noise occurrence in a busy navigation channel to test predictions of compensation, disturbance, and no-modification in calling. We used passive acoustic monitoring to measure the response of Red Drum to vessel noise in the Gulf of Mexico over two spawning seasons (July-October, 2021 and 2022). Our aim was to determine the response of Red Drum to vessel traffic sound exposure levels (SEL) co-occurring over a two-hour crepuscular period during the spawning season. We investigated whether: (1) cumulative exposure of prior vessel noise SEL impacts fish sound production, (2) if fish alter sound production following individual vessel events, and (3) whether fish alter sound production during vessel noise exposure that co-occurs with fish calling. Additionally, (4) we tested if abiotic variables (e.g., water temperature, lunar phase) were predictive of Red Drum calling in conjunction with vessel nose and how this might affect vocal behavior over the spawning season and (5) documented the prevalence of vessel noise cooccurring in the same habit as Red Drum calling.

## Materials and methods

### Data collection and field site

The inlet of Saint Andrew Bay, Panama City Beach Florida (Fig. 1), was monitored via a single passive acoustic monitor (SNAP recorder, 2dB gain, 44.1kHz, Loggerhead Instruments, Sarasota, FL, U.S.A., http://www.loggerhead.com). Recordings took place during the Red Drum spawning seasons (July-October 2021 and 2022; hydrophone sensitivities of recorders ranged from −169.8 to −170.5 dB re 1V/µPa). A chain secured the recorder to a large boulder at a depth of ∼9 m. This site faced the navigation channel in the bay and was observed to be heavily trafficked by both commercial and recreational watercraft. The recorder was set to record 45 second files on a 15% duty cycle. A dive team swapped out the recorder approximately every 45 days throughout the spawning seasons, replacing it with a fresh recorder. A continuous recording, in place of a duty cycle recording, was made during October 2022 via a SNAP recorder installed in the same location (St. Andrew Bay Pass) via scuba as described above. The recorder was left to record 45 second files with no pauses until the end of battery life (5.25 days).

### Abiotic data

Abiotic data were used to assess what environmental factors beyond noise pollution predict Red Drum vocal behavior. For each SNAP deployment an Onset HOBO data logger was deployed at the same location. Hourly temperature readings were averaged by day for a daily value in analysis. Other predictive variables used were daylength, tidal events (time of high tide and low tide, tide differential) for each 24h period (0-24h), time of moonrise, moon phase (illumination %), and time the moon crossed the meridian [31]. If no tidal event occurred in a 24 h period (0-24h calendar day), the next proceeding event was used (24h + difference between events). If two tidal events occurred in the same 24 h period, the highest differential was used.

### Screening recordings

A two-hour crepuscular period (CP) (30 min prior to and 90 min post sunset) was visually and aurally screened with Raven Lite 2.05 software [32] and categorized based on presence/absence of vessel noise and Red Drum calls for each day through both spawning seasons. The CP was chosen for analysis because it encompasses the period when Red Drum spawning activity is most prevalent [17,23]. The CP was analyzed for both duty cycle and continuous recordings with all files bandpass filtered in R [33] using the Seewave package [34]. A Butterworth filter was used to isolate the 0 – 0.6 kHz band. This frequency band was chosen because these frequencies encompass the dominant bandwidth of Red Drum calls [25,26]. We then calculated root mean square (RMS) band sound pressure level (SPL dB re: 1 µPa) for each 15 s interval of the file, which was then averaged for each 45 s sound file. Recordings with noise artifacts from intense weather conditions, such as hurricanes, or the hydrophone being bumped or jostled were not used for analysis. Daily background noise levels were calculated to account for biotic and abiotic variation of background noise. This was done using files from within the CP that contained neither detectable vessel nor Red Drum sounds, though other biotic sounds may still be present in these recordings as they do contribute to natural variation in background noise levels. If full recordings without vessel or Red Drum sounds were not available in the CP, a minimum period of five seconds without vessel noise or Red Drum calling was sampled within a file. If no such period existed within the CP a 5s period was sampled either immediately prior or after the CP to estimate evening background noise levels.

Received sound pressure level (SPL) was used as a measure of fish calling as counting calls was untenable when fish chorused or when vessel noise obscured the soundscape. SPL of each sound source (Red Drum, SPL_fish_; vessel noise, SPL_vessel_) within a file was determined in the following manner. Estimated SPL contribution of the sound source to the total band sound pressure level of the file was calculated using the power summation equation [35] and subtracting the estimated contribution of evening background noise from the total band RMS sound pressure level of the file. Vessel noise SPLs may also contain Red Drum sound, but it was not possible to directly count calls or estimate the contribution of Red Drum calls when vessel noise obscures potential fish calls. We explored this possibility below (*see Immediate impacts of vessel noise during fish calling)*. In a few cases, total band SPLs were lower than estimated background noise levels (likely due to limits in precision of background noise estimates). In these few cases, SPLs of vessels or fish were estimated as zero. We estimated total contribution of fish calls and vessel noise of each CP as a sound exposure level (SEL dB re: 1μPa^2^ ⋅s). SEL of fish calls for each CP was estimated as the following:

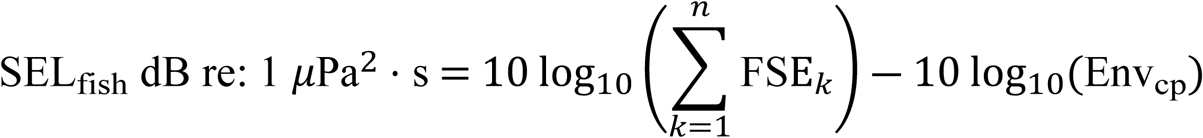

The number of files with fish calling is represented by *n. FSE* is the sound exposure of a single 45 s file, estimated by squared SPL_fish_ (µPa^2^). *Env_cp_* is the number of files without vessel noise present for the whole CP (*Env_cp_*<u>≤</u> 24). Thus, *Env_cp_* accounts for the amount of time (CP or portion of CP without vessel noise) for which SEL_fish_ is estimated. Thus, SEL_fish_ accounts for sustained levels of FSE across the CP. For example, a single instance of fish calling at 129 dB re: 1µPa^2^ out of 24 files would be approximately equal to 24 instances of a fish calling at 101 dB re: 1µPa^2^, ∼ 115 dB re: 1µPa^2^ ⋅s.

Vessel noise SEL was calculated in a similar manner:

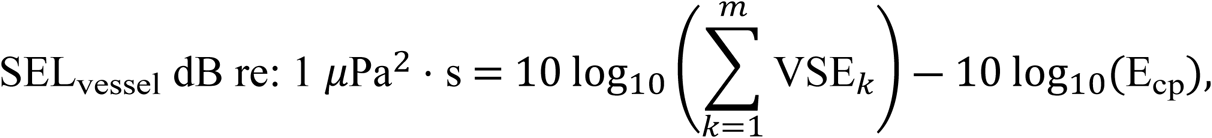

where VSE is the SPL in µPa^2^ of a sound file with vessel noise present, *m* is the total number of files with vessel noise for the whole CP on that date, and *E_cp_*is the total number of files of the CP on that date.

Files from continuous recordings were visually and aurally screened and categorized as above. When vessel noise was detected, recordings were band pass filtered (0-600 Hz) as described above and the band pressure of filtered files was estimated from RMS amplitudes measured in Audacity software to determine SL within specific periods of time relative to vessel noise occurrence (see section Fish calling during and immediately following individual vessel noise events, below).

### Relationship of fish calling to vessel noise and other abiotic variables

We tested the combined influences of vessel noise and potential environmental predictor variables on fish calling SEL over the CP among dates. To determine which predictor variables had the greatest influence on fish calling SEL by evening, complex Generalized Least Square (GLS) models were constructed in R package nlme [36]. GLS models are ideal for time series data where autocorrelation between residuals is likely to occur. Initial models included all abiotic measurements (vessel noise SEL, water temperature, daylength, time of high tide, time of low tide, tide differential, moon phase, moonrise time, and time of moon passing the meridian). Date was also included as a predictor and coded with consecutive integers beginning with 1 for 1 July. Additionally, an interaction of date and daylength was included as a predictor, as daylength shortened throughout the months inversely with increasing date. Red Drum spawning season ends prior to the shortest day of the year. These models were simplified in a backwards stepwise fashion to obtain the combination of variables which produced the lowest Aikake Information Criterion value corrected for small sample size (AICc). From the best fit model, predictor variables were removed if their inclusion raised the model < 2 AICc units [37]. For these models we also tested for serial autocorrelation [38] based on the expectation that fish calls may cluster among consecutive days. We conducted backwards stepwise model selection (as described above) for the first (AR1) and second order (AR2) autoregressive models and a non-autocorrelation (non-AR) model. We then selected the best model among these three models and three null models (non-AR, AR1, and AR2) based on AICc.

### Prior exposure impact on Red Drum calling

To test whether prior vessel noise exposure affects fish calling later in the evening, we used GLS models to determine whether fish calling SEL in the final portion of the CP (75-90 min.) is predicted by prior (30 min. pre-sunset to 75 min. post-sunset) vessel noise exposure. Although Red Drum calling may be less intense 75 min. post-sunset, we chose to examine this period in order to determine if prior vessel noise influences fish calling. Thus, our analysis compared fish calling among days at the same time point relative to sunset. In addition to prior vessel noise, the same abiotic variables used to test fish calling SEL over the CP were used in initial GLS models that were simplified step-wise (as described above). Data from CPs without fish calling from 75-90 min were not included in this analysis as it was unknown if fish were still present in the recording area.

### Fish calling during vessel noise compared to periods without vessel noise

Vessel noise often obscured the soundscape to a degree that visual (via spectrograms) and aural observations of fish calling were not possible (Fig. 2). During intense vessel noise, fish may reduce calling because of behavioral disturbance. However, direct assessment of fish calling during intense vessel noise is not possible because this noise would obscure potential calls. If fish maintain the same call amplitude during vessel noise as in quiet periods, the resulting SEL during vessel noise should be higher than during periods with fish calls alone, as the non-coherent combination of fish sounds and vessel noise would be additive. Observations with vessel noise and potential fish calling (VN/FC) at a lower SEL than prior fish calling by itself would only be expected if fish reduced calling amplitude during vessel noise, by either lessening calling or moving away from our study site, as predicted by the disturbance hypothesis. However, high amplitude vessel noise (relative to SEL of fish calling alone) could result in vessel noise recordings with higher SELs than periods with fish calling even if fish reduce calling during vessel noise. Thus, to examine the potential of fish modulating call amplitude during vessel noise, we focused our analysis on evenings in which fish calling SEL was equal or greater than the median SEL observed during vessel noise. We analyzed 2021 and 2022 separately and used a Wilcoxon sign rank test to determine if SEL during vessel noise periods differed from fish calling SEL measured on the same evenings.

**Fig 2.**
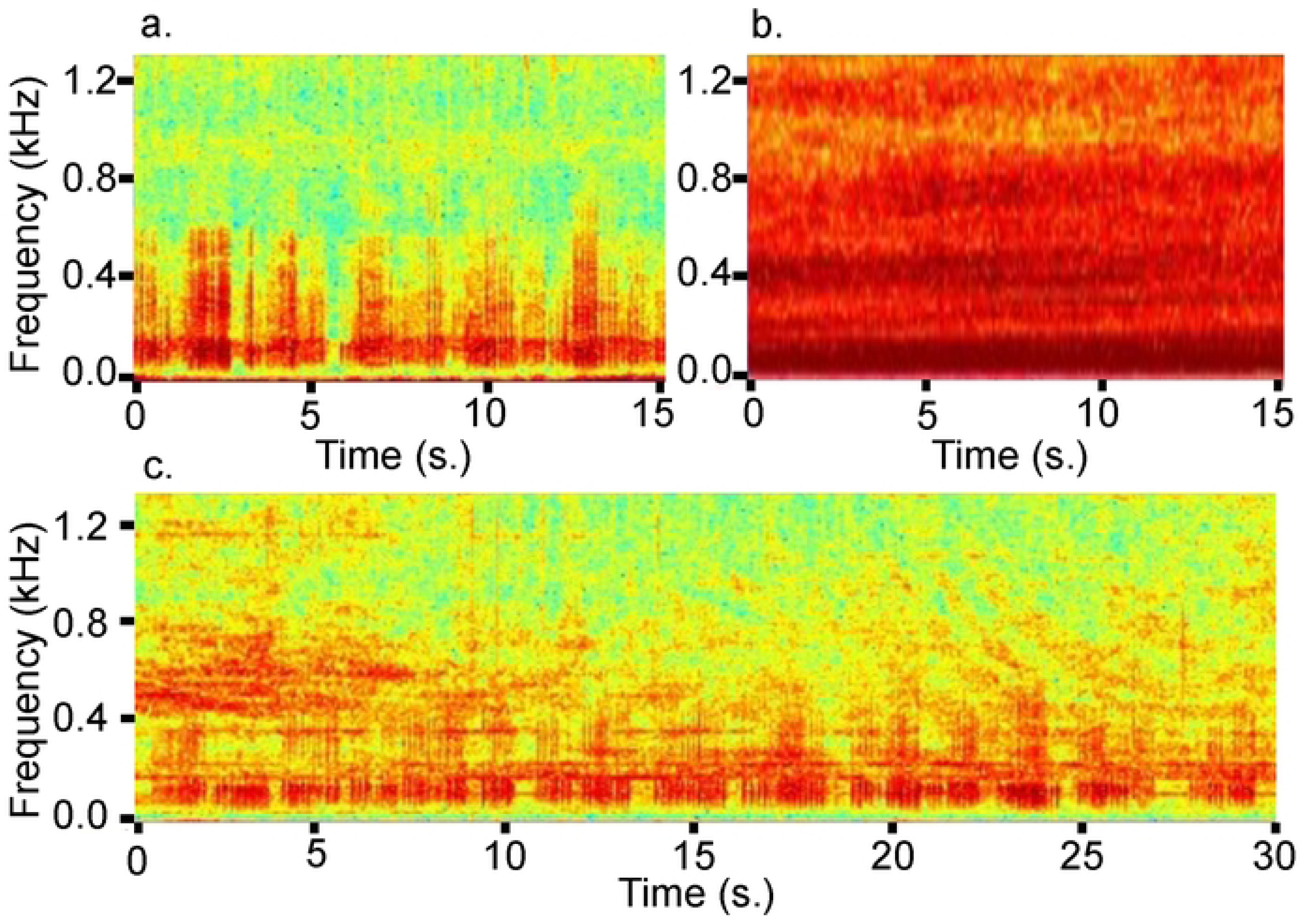
Spectrograms. Spectrograms showing field recordings (0-1kHz) at Saint Andrew Bay, FL. (**a**) Red Drum calls in the 0.1-0.5 kHz range. (**b**) High-intensity vessel noise. (**c**) Red Drum calls co-occurring with characteristic low-frequency vessel noise. Of note, high intensity vessel noise (b) encompasses a large percentage of the soundscape, which likely obscures Red Drum calls from visual and aural observation.

### Fish calling during and immediately following individual vessel noise events

We used continuous recordings (Oct. 2022) to examine the acute response of fish calling during and immediately following individual vessel noise and potential fish calling (VN/FC) events. To investigate the potential impacts of vessel noise on fish calling during and immediately after, we examined SELs of individual periods with VN/FC and periods of only fish calling immediately prior and following. We identified VN/FC events from continuous recordings and measured fish calling (F1) SEL before VN/FC (one min), SEL during VN/FC, and fish calling (F2) SEL immediately afterward (one min). VN/FC events were considered as the entire period of continuous recording in which vessel noise was detectable visually and aurally. Each VN/FC event was divided into 20% increments (V1-V5) with SEL taken for each period. We restricted observations to VN/FC events with >1 min vessel free immediately prior to and following each event. Each vessel-free, fish calling period could not be used as prior or following for multiple VN/FC events. Additionally, we calculated hypothetical vessel noise amplitudes based on potential fish call levels and graphed these alongside the original recording values. These hypothetical vessel amplitudes were used to illustrate if hypothetical behavioral shifts could occur based on the sound levels taken in the original recordings.

### Spatial and Temporal Overlap of Vessel Noise and Red Drum Choruses

For each CP we tallied the total number of recordings with either vessel noise or Red Drum calls without any vessel noise present. From these values, for each daily CP we calculated ‘fish calling tendency’ (fct), ‘vessel noise portion’ (vnp), ‘noise-free fish calling portion’ (n-ffcp), and the noise free portion (nfp), as the following:

*fct* = (no. files with fish calls but without vessel noise) / (no. files without vessel noise) *100
*vnp* = (no. files with vessel noise) / (total no. of files in CP)*100
*n-ffcp* = (no. files with fish calls but without vessel noise) / (total no. of files in CP) *100
*nfp* = (no. files without vessel noise) / (total no. of files in CP)*100

We expected the fct to reflect calling propensity changes that parallel seasonal changes in spawning activity. Values of vnp likely follow seasonal patterns of vessel traffic in the area and the n-ffcp reflects the amount of the CP occupied by fish calls without vessel noise as a combination of fct and the size of the nfp. For both seasons, we summarized the monthly mean, median, quartiles, interquartile range, and outliers of the fct, vnp, n-ffcp, and nfp.

## Results

### Relationship of fish calling and vessel exposure with abiotic variables

During CPs in both spawning seasons our recordings captured Red Drum calls (Fig. 2a), vessel noise (Fig. 2b), and their co-occurrence (Fig. 2c). In 2021, Red Drum total crepuscular calling SEL by date was negatively correlated with evening vessel SEL, after inclusion of abiotic variables that were most strongly predictive (Table 1, Fig. 3). The interaction of date and daylength was the most supported predictor in the model (Table 1). This interaction predicts an increase in calling associated with the reduction of daylength, but with calling peaking (highest evening SEL value, September 15, 2021, October 4, 2022) prior to the shortest day of the year of our study period. Additionally, calling was positively correlated with moon illumination and tidal differential during the 2021 spawning season. In 2022 there was also an interactive effect between date and daylength with these acting as the sole predictor of total crepuscular calling SEL by date (Table 1, Fig. 3). Notably, 2022 recording period was shorter than 2021 with the last half of the month of October not being collected. Additionally, the auto regressive model was supported in 2022 but not for 2021. This indicates Red Drum calling SEL was more clustered over successive days in 2022.

**Fig 3.**
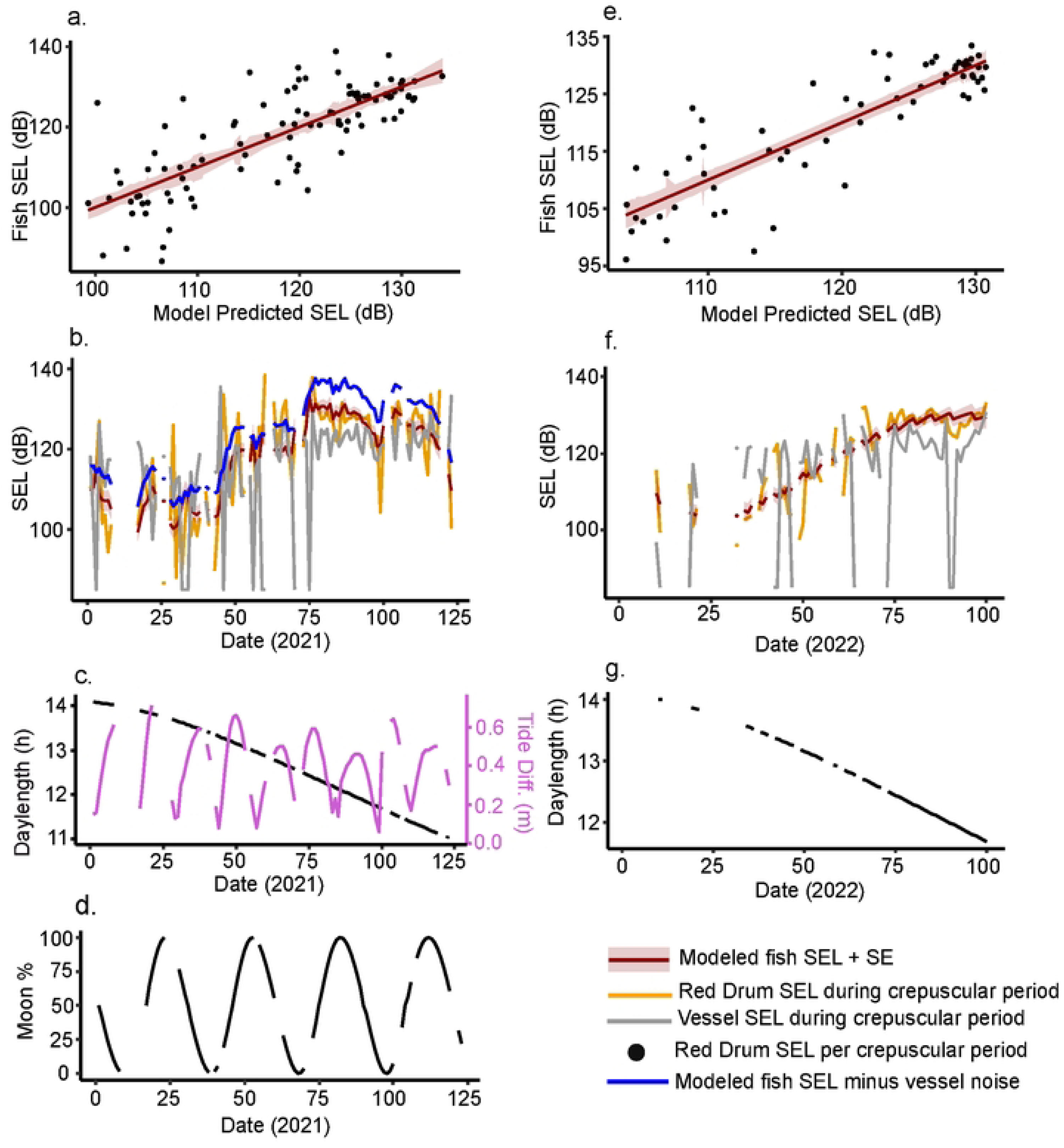
Nightly Calling. Red Drum call sound exposure level (SEL) (dB re: 1µPa^2^·s) over a daily two-hour crepuscular period with GLS models used to determine association with vessel noise and other abiotic variables. (**a**) Best-fit model for 2021 spawning season included vessel SEL, moon phase, tidal differential, and an interaction of date and daylength. (**b**) Fish call SEL (gold) and vessel (grey) over the 2021 spawning season along with the fitted line (red) and the model without vessel as a predictor (blue). (**c**) Daylength (black) and tidal differential (purple) over 2021 spawning season. (**d**) Moon phase (illumination) over 2021 spawning season. (**e**) Best-fit model for 2022 included an interaction of date and daylength but did not include vessel SEL. (**f**) Fish call SEL (gold) and vessel (grey) over the 2022 spawning season along with the fitted line (red). (**g**) Daylength over 2022 spawning season. Note for 2021, the model without vessel noise included would predict elevated levels for fish SEL (blue line).

**Table 1.**
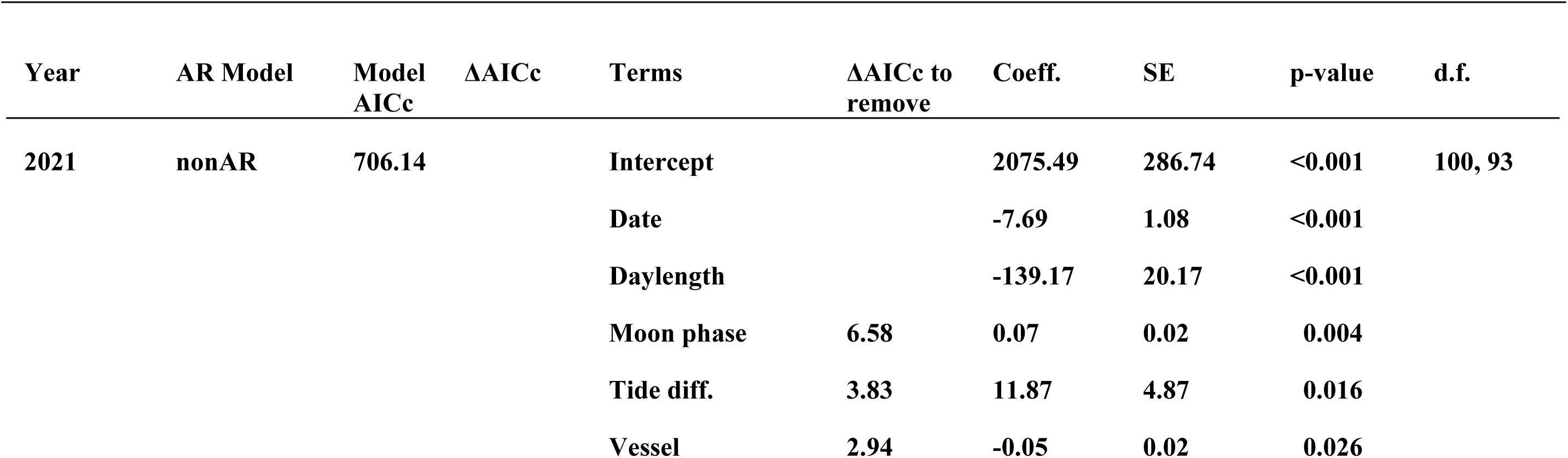
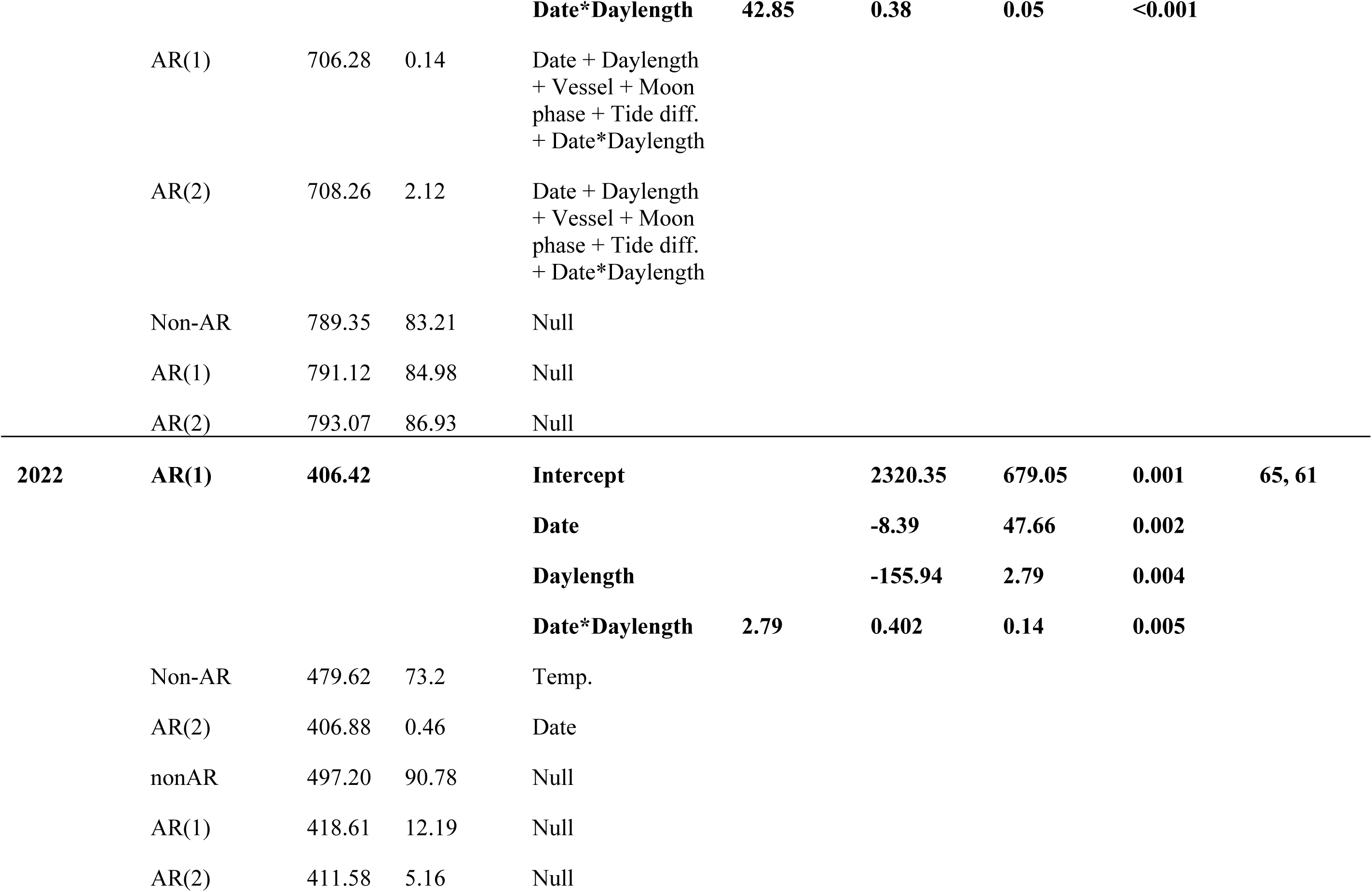
GLS Models for crepuscular periods over two spawning seasons (2021-2022)

### Prior exposure impact on Red Drum calling

In accordance with expectations of the disturbance hypothesis, prior sound exposure to vessel noise exposure during the first 105 min. of the CP predicted a reduced fish calling amplitude during the final 15 min. of the CP for both 2021 and 2022 (Fig. 4, Table 2). This indicates that the level of fish calling observed in the final 15 mins of the CP was negatively correlated with the SEL of vessel noise during the 105 min. prior. Additionally, day length correlated negatively with Red Drum SEL during both spawning seasons. Moon phase and temperature were both positively correlated with fish SEL in 2022 only (Table 2, Fig. 4f, g).

**Fig 4.**
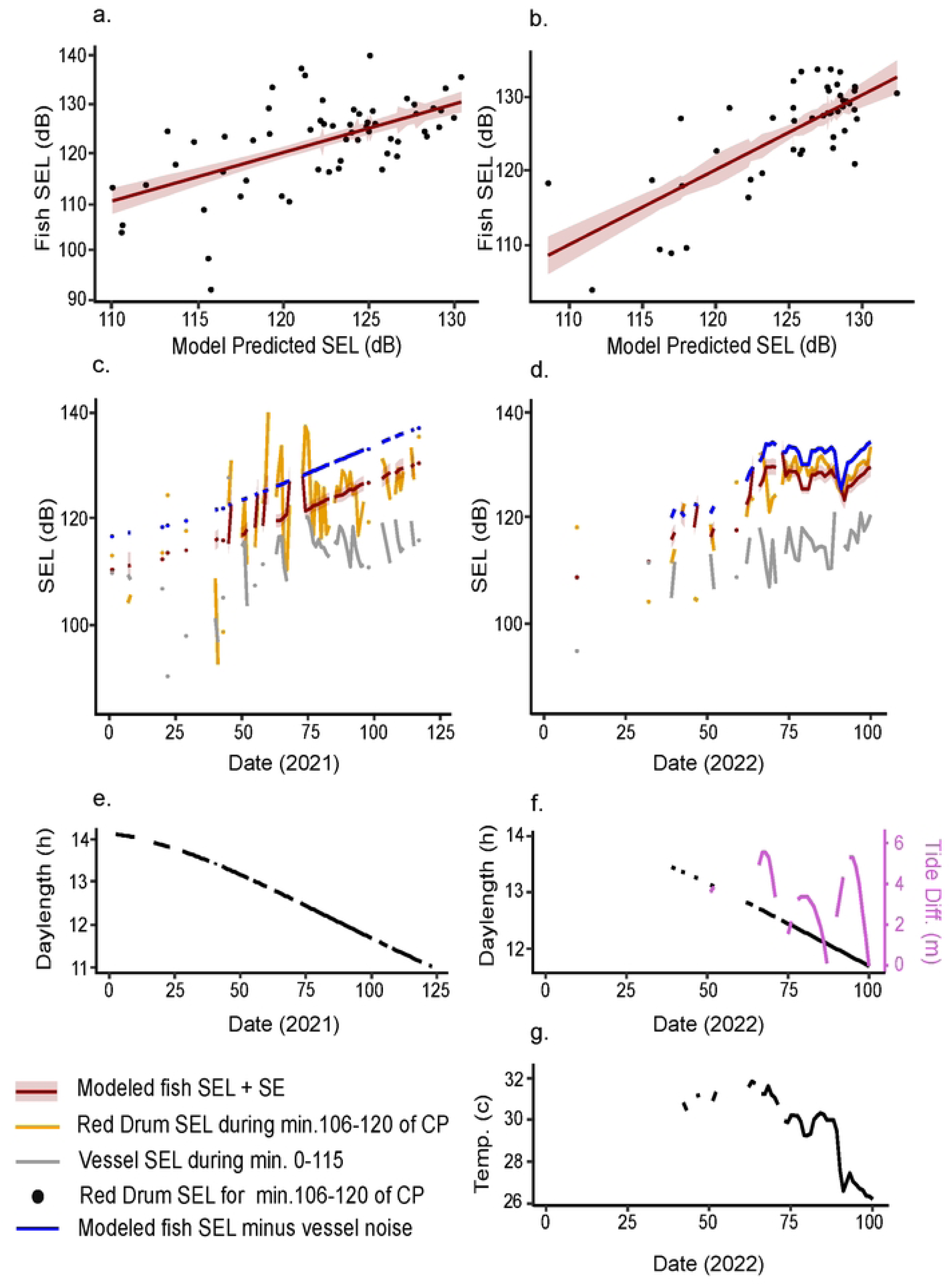
Prior Noise Impact. Prior vessel noise predicts quieter Red Drum calling sound exposure level (dB re: 1µPa^2^·s) (SEL). Best fitted models for Red Drum calling amplitude over a 15 min period, 90 min after sunset, predicted by daily abiotic variables in the 105 min prior for 2021 (**a**) and 2022 (**b**). For 2021 prior vessel SEL and daylength were both included in the best fit model. For 2022 prior vessel SEL, daylength, moon phase, and temperature were all included in the best fit model. (**c-d**) Vessel SEL (grey) in the initial 105 min of the crepuscular period (CP), Red Drum call SEL (gold) for the final 15 min of the CP and the fitted line (red) over both 2021 and 2022 spawning seasons along with the model minus vessel noise (blue). (**e-g**) Abiotic predictors for both spawning seasons; daylength (**e**) 2021, daylength, moon phase (purple), and temp. (**f, g**) 2022. Note that for both 2021 and 2022, the model without vessel noise included predicts elevated fish SEL (blue line).

**Table 2.**
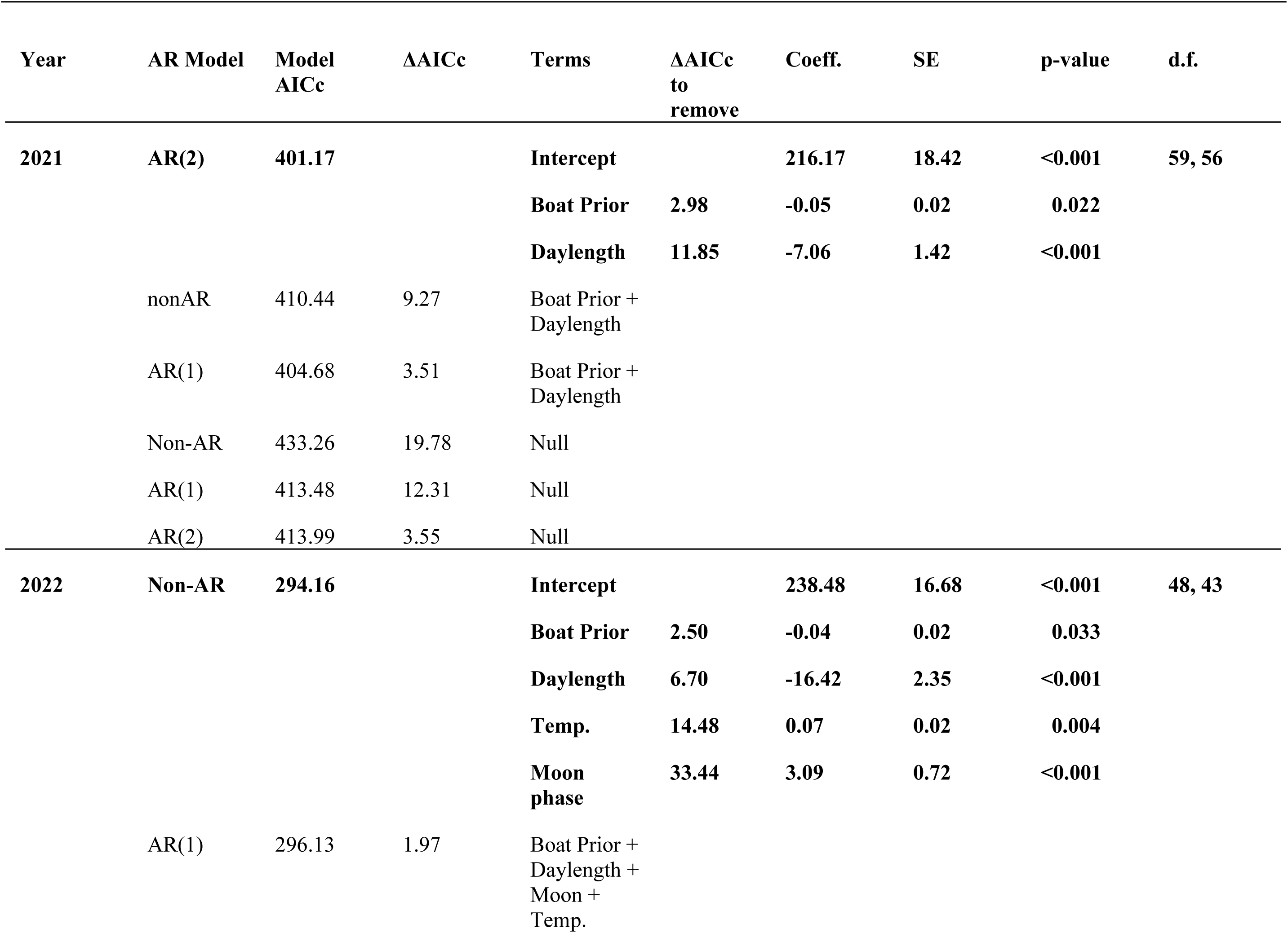
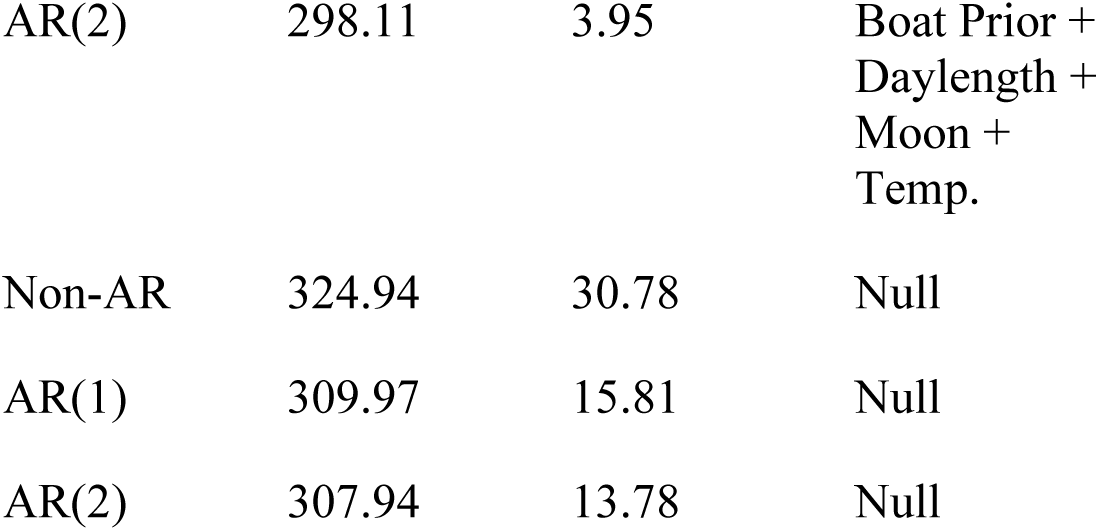
GLS models for final 15 min of crepuscular period over two spawning seasons (2021-2022)

### Fish calling during vessel noise compared to periods without vessel noise

In both 2021 and 2022, when fish calling SEL was low to moderate, SEL during vessel noise periods regularly exceeded fish calling SEL (Fig. 5). In these instances, it is not possible to determine if fish calling SEL changed during vessel noise because there is a greater likelihood of vessel noise amplitude exceeding the amplitude of fish calling alone. In 2021, on evenings when fish calling SEL was > 125.1 dB (median 2021 vessel noise amplitude), SEL during vessel noise periods did not differ (n=36, p = 0.499, Wilcoxon sign rank test) (Fig. 5A). In 2022, when fish calling SEL was > 124.6 dB (median 2022 vessel noise amplitude), SEL during vessel noise periods was lower than fish calling SEL (2.5 dB median difference, 0.5-4.9 dB quartile 1-quartile 3) measured on the same evening (n = 35, p <0.001, Wilcoxon sign rank test) (Fig. 5B). Lower SEL values during vessel noise periods is consistent with the expectations of the disturbance hypothesis.

**Fig 5.**
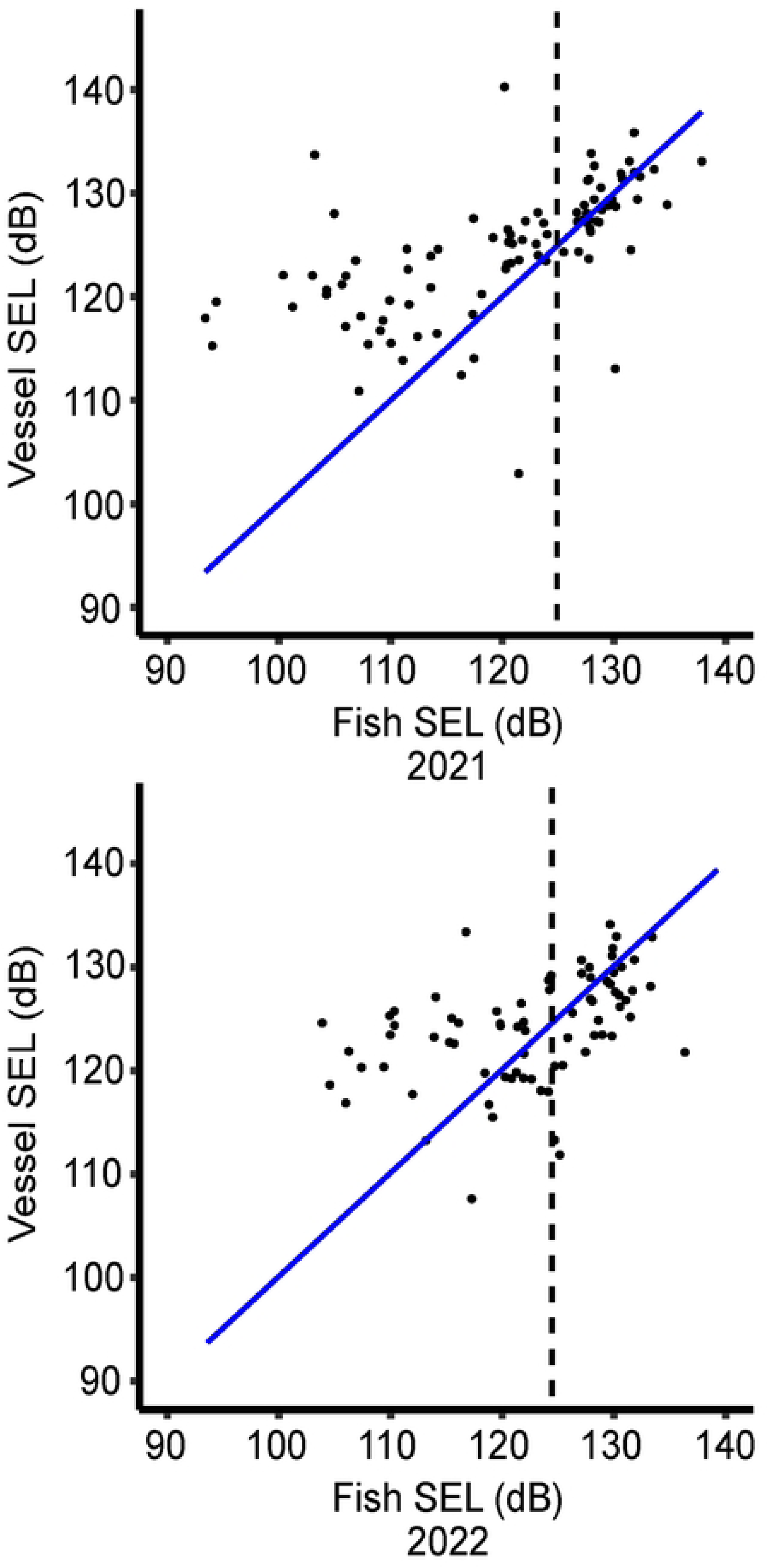
Possible Fish SEL. Sound exposure level (SEL) (dB re: 1µPa2·s) of vessel noise with potential fish calling vs. SEL of recordings with fish only in 2021 (a) and 2022 (b). The blue, 1:1 line represents the minimum possible SEL during periods of vessel noise if fish calling remained unchanged. Points below the blue line are consistent with predictions of the disturbance hypothesis. Points above the 1:1 line are ambiguous, and it is impossible to assess what contribution fish calling may have made to these data points as high intensity vessel noise will not be significantly altered by background noise. The dotted vertical lines indicate the median SEL during vessel noise periods in 2021 and 2022, respectively. Differences between SEL during vessel noise periods and SEL of fish calling alone were tested with a Wilcoxon sign rank test when fish calling was > median SEL of vessel noise periods (i.e., to the right of the dotted line).

### Fish calling during and immediately following individual vessel noise events

In observations from continuous recordings (Fig. 6, Fig S1), SEL during vessel noise periods was lower at times than fish calling SEL immediately prior (F1) in 13 of 23 cases. Because the contribution of vessel noise to the combined VN/FC SEL is not known, it is not possible to determine if fish modulate their calling amplitude in situations in which VN/FC SEL is higher than prior fish calling SEL (F1). It is possible, however, to estimate what the contribution of vessel noise to the combined VN/FC SEL would be if fish continued to call at the same amplitude of prior calling (F1). In seven of the 10 instances in which the combined VN/FC SEL exceeded F1 amplitude, vessel noise would be less than 115 dB for at least a portion of the measured period, which is relatively low (observed <7% of vessel recordings on nights when Red Dum were not observed calling in the study) (Fig 6, Fig S1). Thus, vessel noise would need to be relatively low during these instances to observe these combined SELs if fish calling remained at F1 levels. It is also possible to estimate what the combined SEL of fish vessel noise and fish calling would be if fish compensated with an increase in calling amplitude of 5 dB during vessel noise. There were only six of 23 cases in which a 5 dB increase in fish calling would be possible during vessel noise (Fig. 6b, Fig. S1m,n,p,q,s).

**Fig 6.**
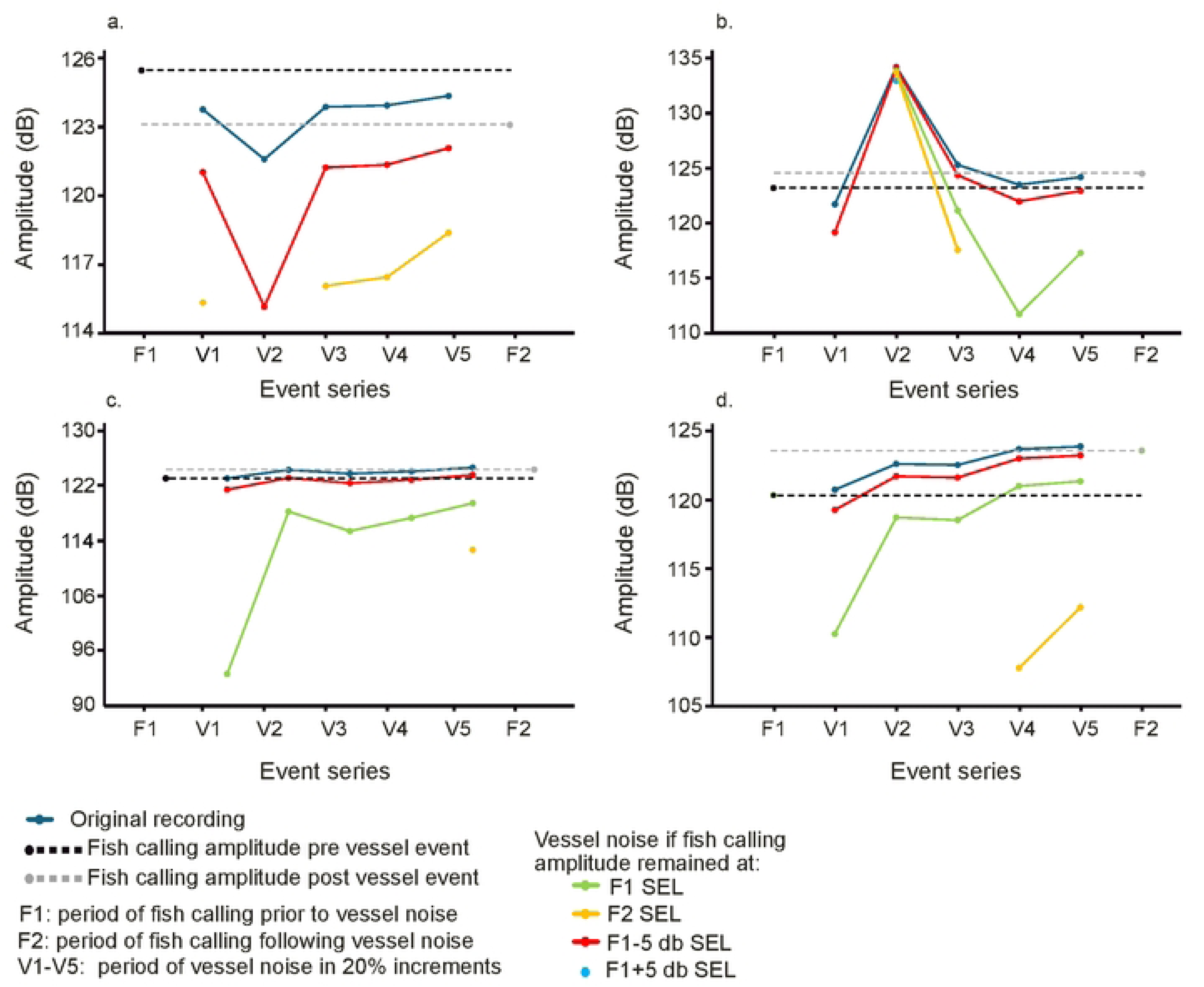
Continuous Recordings. Sound exposure levels (SELs) (dB re: 1µPa^2^·s) from continuous recordings of calling fish immediately prior (F1, fish SEL prior to vessel noise), during (V1-V5, five 20% increment measurements of vessel noise), and after vessel noise occurrence (F2, fish SEL after vessel noise is no longer present). Seven discrete SEL values are depicted (see Methods for additional details) and lines between successive points are shown for ease of comparison. Theoretical vessel SEL values of five separate hypothetical fish calling SELs that would combine to result in the observed SEL during this period (V1-V5). Missing lines and points indicate cases in which these theoretical fish calling SELs could not combine with vessel noise to form the observed SELs. (**a**) Potential calling disturbance in which SEL during vessel noise is lower than fish calling SEL prior to vessel noise occurrence and remains lower following vessel noise. (**b**) Evidence of potential calling disturbance during portions of vessel noise. SEL prior (at V1) to the peak of vessel noise SEL is lower than the initial calling SEL. Higher SEL values from V2-V4 could occur if vessel noise amplitude exceeds prior calling amplitude or if fish increase calling amplitude during this portion of vessel noise. (**c-d**) Instances in which SEL during vessel noise exceeded prior calling SELs. In both examples (**c-d**), post-vessel fish SELs were higher than pre-vessel levels. In these two examples (**c-d**), the SELs during vessel noise may be higher than preceding calling SELs either because vessel noise was higher amplitude or because fish calling co-occurred and combined to increase the observed SEL during vessel noise.

### Spatial and temporal overlap of vessel noise and Red Drum choruses

In Saint Andrew Bay, the monthly mean of recordings that contain vessel noise (vnp) in the CP ranged from 14% in August (Fig. 7c.) to 45.1% in October (Fig. 7g) in 2021 and 31.3% in September (Fig. 7f.) to 53.4% in July (Fig. 7b) in 2022. Two days in 2021, October 9-10, contained vessel noise in 100% of the recordings during the CP. The highest percentage of the CP to display vessel noise in 2022 was 87.5% on July 23. In both spawning seasons, Red Drum fish calling tendency (fct), the percentage of the noise free portion (nfp) (non-vessel noise portion of the CP) to contain fish calling, was higher in late-summer and early autumn at over 90% in September-October (Fig. 7e-h). Note that this period of high fct largely corresponds with the period for which high fish calling SELs were observed, days 75-100 (13 September – 8 October) in both years.

**Fig 7.**
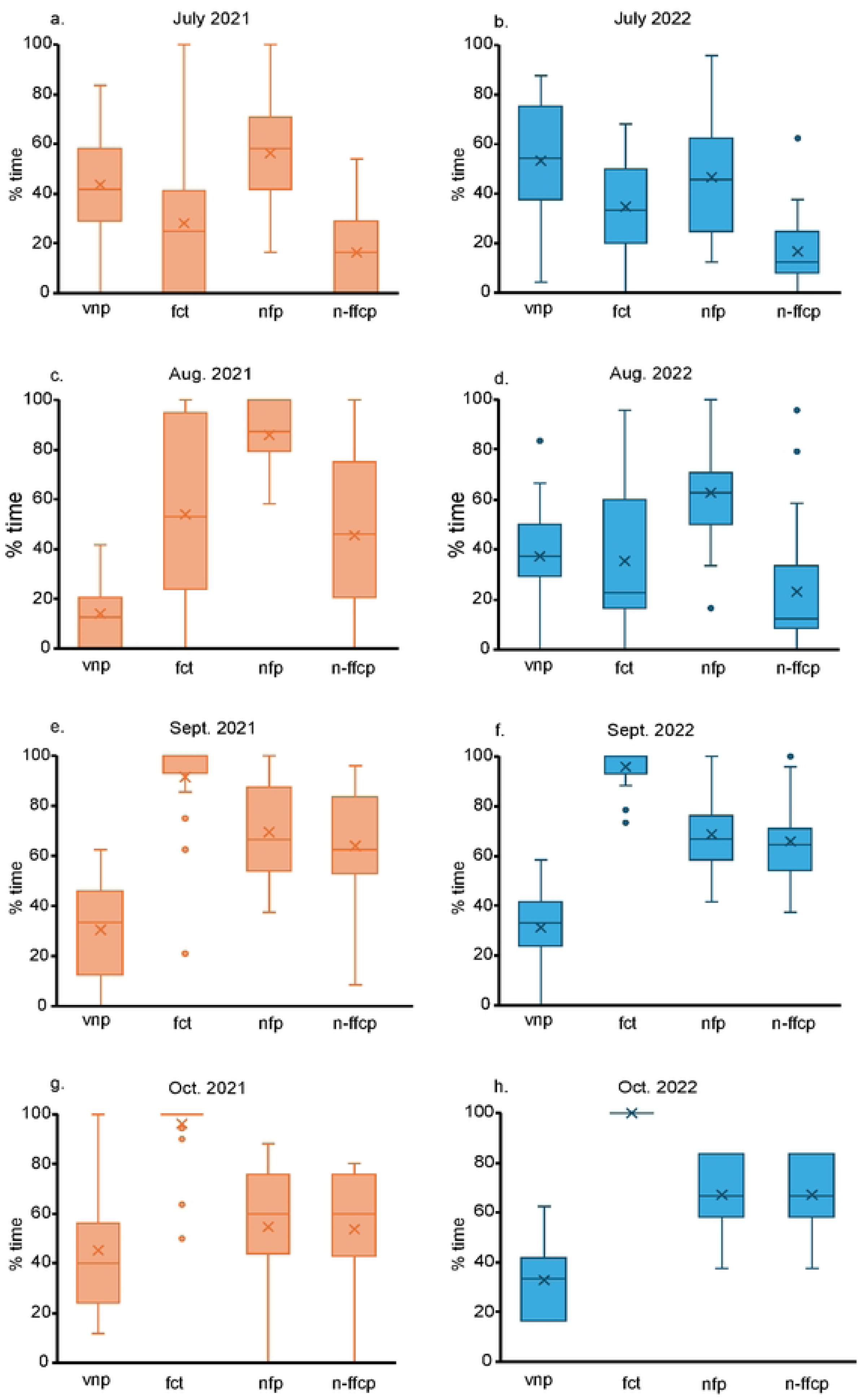
Temporal overlap. Percentage of crepuscular period sound files by month in 2021 and 2022 spawning seasons that contain vessel noise (vessel noise portion, *vnp*), fish calling tendency (*fct*), lack vessel noise (noise free portion, *nfp*), and contain fish calling (noise-free fish calling portion, *n-ffcp*). Fish calling tendency (*fct*) is the percentage of crepuscular *nfp* sound files containing fish calls. Boxes indicate 25^th^-75^th^ percentile, line indicates the median, x indicates the mean, whiskers encompass data range excluding outliers, dots are outliers.

## Discussion

Our findings indicate potential disturbance of Red Drum spawning aggregation calls during vessel noise. The disturbance hypothesis was supported, with data reflecting lower total crepuscular fish call SEL in association with total crepuscular vessel noise in 2021. Evidence of disturbance from total crepuscular fish call SEL in association with vessel noise was not observed in 2022, but the recording period was shorter for 2022 and did not capture behavior of Red Drum as late into the season as 2021.

SEL of vessel noise 105 min prior to the end of the CP was shown to have a negative correlation with SL of fish in the final 15 min. of the CP. The results, in support of disturbance hypothesis, were observed in both 2021 and 2022, when lower fish call SEL was associated with higher SELs of anthropogenic noise in the preceding period. This analysis shows the impact of vessel noise is not just an immediate issue, as prior vessel noise exposure can also affect fish behavior. Lower call amplitudes observed in this analysis at the end of the CP indicate that Red Drum do not acclimate to noise exposure over the CP. This acute response in call amplitude could be due to reductions in the number of calls or duration of calling by individual fish, or fish may vacate habitat with increased vessel noise. This contrasts with research in the Gulf of Mexico off Florida which found no evidence of vocal modification in response to ferry noise in a closely related species, Atlantic Croaker (*Micropogonias undulatus*) [41].

Notably, without visual confirmation of fish proximity, we are unable to determine if fish remain near the recorder. However, we do not believe our findings are due to fish vacating habitat alone for multiple reasons. Firstly, Red Drum do not appear to permanently vacate the area following noise exposure as Red Drum sounds were recorded over the CP throughout the spawning season, over consecutive nights, even though vessel noise was ubiquitous. Interestingly, the sciaenid Brown Meagre (*Sciaena umbra*) was found to occur in noisy habitats near Venice Lagoon, and less so in quieter habitats which indicates that factors other than vessel noise likely exert a stronger influence on habitat choice [42]. Additionally, fish may have other means of signaling beside acoustic communication that they employ when background noise becomes too intense. Two gobiids were observed in lab settings to reduce spawning sounds but continued with visual communication in the presence of increased background noise [7]. These gobiid fishes still spawned in the presence of anthropogenic noise, though were forced to deviate from their typical behaviors. Perhaps behavioral plasticity exists in other species, though in turbid environments, such as those often inhabited by nearshore fishes of the Gulf of Mexico, visual communication would likely be less effective.

On evenings with the highest fish calling SEL, we observed evidence that fish either decrease calling SEL, fewer total fish call, or fish move away from the recording area during vessel noise, consistent with predictions of the disturbance hypothesis. In 2022, when fish calling SEL was as high or greater than the median SEL of vessel noise periods, SEL during vessel noise tended to be lower than fish calling SEL from the same evening. This indicates that fish call amplitude may be diminished in the presence of vessel noise or that fish move away from the recorder during times when vessel noise is present. If fish were to maintain or increase call amplitude, recordings with vessel noise over the same evening should exceed fish calling SEL alone as the two signals should be additive, resulting in a higher amplitude than a single signal [48]. This predicted disturbance response can only be tested when fish calling SEL is greater or equal to vessel SEL because an effect cannot be detected when fish call at amplitudes lower than the concurrent vessel noise. However, we are unable to determine if fish calling SEL is modified when SEL during vessel noise presence exceeds SEL of fish calling alone because this could result either when vessel noise amplitude exceeds fish calling SEL or when fish calling and vessel noise combined result in a higher amplitude than fish calling alone. In 2021, when fish calling SEL was greater or equal to the median SEL of vessel noise periods, SELs of fish calling were not found to differ from SELs of vessel noise periods; i.e., there was neither evidence of a reduction (disturbance) in fish calling during vessel noise nor evidence of an increase in fish calling during vessel noise (compensation). Notably, SEL during vessel noise was slightly higher in 2021 compared to 2022 (median 125.1 vs. 124.6 dB) and fish calling SEL in 2021 was slightly lower compared to 2022 (median 121.5 vs. 124.1 dB). Thus, there may be a greater opportunity to test predictions of the disturbance hypothesis when fish calling amplitudes are higher than received vessel noise amplitudes, which may have been the case in 2022. Also, it should be noted Red Drum may respond differently between high or low intensity vessel noise or when calling at lower sound levels, i.e., compensate under low levels of vessel noise and act disturbed under higher vessel noise.

Reduction in calling amplitude during vessel noise was observed in some cases, 13 of 23 cases from continuous recordings. Reductions in calling during vessel noise predicted by a disturbance hypothesis are reported for several sciaenid fishes: Meagre (*Argrosomus regius*) calling during vessel noise in the Tagus estuary of Portugal [8], and *Pogonias courbina*, decreased call rate during vessel noise in Argentina [45]. Meagre were reported to reduce spawning choruses after ferry boat passage, while Weakfish (*Cynoscion regalis*) displayed an escape response without a change in call rate in the same study [40]. These findings also align with preliminary analysis of a subset of our data set showing disruption of Red Drum calling by vessel noise [46]. The results of our study, which expands the scope of preliminary work, and other recent investigations indicate that sciaenid behavioral responses to vessel noise may vary by location and among species due to population differences, environmental factors, or a combination of both [47]. In the 10 of 23 cases in which SEL of vessel noise periods exceeded prior fish SEL, however, it is not possible to determine if fish modulated their calling amplitude because it is possible that the received amplitude of vessel noise exceeded prior fish calling amplitude. Notably, in most cases when vessel noise periods exceeded prior fish calling SEL, if fish were to have called at the same prior calling amplitude, vessel noise would be relatively low amplitude relative to most of our observations. Thus, it is possible that fish reduced their amplitude but the combination of vessel noise and fish calling exceeded prior fish calling SEL. It is also conceivable that fish may respond differently based on the intensity of background noise, e.g., maintaining or even increasing calling amplitude when vessel noise is low (no response or compensation) and reducing calling when vessel noise is more intense (disturbance). For future studies, an experimental approach with a known vessel noise source that passes repeatedly over a study area with the same path and speed that could be measured in the absence of fish calling (e.g., earlier in the day before chorusing occurs) and during choruses could test these responses but would also require that other vessel noises are not present, which would be a challenge at this study site.

Factors that should be considered when interpreting received fish sound levels include the water depth, bathymetry, and reflection from surrounding objects. In our study, these factors were relatively constant; the recorder was at a fixed location, and tidal range is relatively small at St. Andrew Bay. Red Drum sound received levels will also be influenced by source level, position and distance from the hydrophone. Our analyses assumed that Red Drum were randomly dispersed around the hydrophone and thus over the course of each study night, the chance of detection and loss from attenuation with distance was not correlated with vessel noise. We routinely observed adult Red Drum near the recorder during dives to deploy and service the recorder. We were not able to directly estimate signal attenuation and detection range in our study, which would provide an estimate of received level variation associated with fish movement away from the hydrophone when call rate is constant. Because the study site is in relatively shallow water, attenuation from geometric spreading (ignoring absorption) is expected to fall between cylindrical (SL = RL + 10·log_10_R) and spherical spreading (SL = RL + 20·log_10_R), where SL = source level, RL = received level, and R = source distance from the hydrophone. Using our minimum detectable received level of Red Drum choruses (108 dB) and a potential high amplitude source level of 150 dB [22], there is wide variation in the detection radius predicted by spherical spreading (126 m) and cylindrical spreading models (15.8 km).

During most of the spawning season, received SELs varied more modestly (10-20 dB). Reductions in received SEL from calling fish in the present study following periods of vessel noise could result from fish moving further away from the recorder. A 10 dB reduction in received level would result if fish were present at the source and subsequently moved away (3.2 m under spherical spreading or 10 m under cylindrical spreading). Fish initially detected at a moderate distance from the hydrophone, however, would need to move much further away from the source to result in a 10 dB reduction. Future studies with a synchronized hydrophone array could be used to reduce error associated with source (fish and vessel sound) position for Red Drum and potentially determine if calling fish move away from noise sources or simply alter call rate [40].

Analyses used in our study assumed that sound amplitudes combined from multiple sources (individual fish, vessels, and background noise) did not combine constructively or destructively. This was based on the observation that Red Drum calls are trains of short duration pulses separated by a silent interval, rather than tonal signals. Thus, we assumed these sources combined additively as predicted for non-coherent signals [48], as has been assumed in other field studies that examined fish sound production [49, 50]. Use of a hydrophone array in future studies may shed light on how fish and vessel sounds combine at different locations within the estuary to determine the validity of these assumptions.

In our study, we did not determine vessel noise source level and location. Source levels and frequencies vary widely depending on vessel type and operation [10]. Our aim was not to determine noise level from individual vessels, but rather to determine the received sound level at the recorder location from all vessel noise sources regardless of the number of vessels, operation, and source location. Our recordings represent what Red Drum may be exposed to in the study site by observing noise pollution present regardless of source. Future work with non-duty cycled recordings and hydrophone arrays is needed to better assess how source levels from individual vessels influence spatial patterns of Red Drum calling. This would allow for a more comprehensive understanding of the spatial patterns of vessel noise and fish calling and could potentially determine if calling fish move in response to vessel noise.

Though we observed reduced Red Drum call SEL during and in association with vessel noise, this pollution source is a relatively recent potential selection pressure for the evolution of sound production behavior in fishes and evidence exists that some sciaenids, along with other taxa, do not avoid noisy waterways or alter calling behavior during vessel noise [12, 42]. Brown Meagre showed an increase in call rate after repeated vessel noise exposure [5, 51]. Plainfin Midshipman (*Porichthys notatus*) and Oyster Toadfish (*Opsanus tau*), both batrachoidids, increase their call amplitude when artificial noise is introduced to calling males [43, 44]. In addition to increasing amplitude, Plainfin midshipman also lower their call frequency, potentially to avoid masking by noise pollution [44]. Freshwater Drum (*Aplodinotus grunniens*) may also vary their peak call frequency in the presence of vessel noise when calling at high amplitudes [52]. Though a frequency shift was not measured in our study, if Red Drum do shift call frequency, it would be highly unlikely that such a shift would occur outside the 0–600 Hz band observed. Additionally, there exists the potential that fish may have shifted sound production to another time outside the CP. Though in situ observations have previously shown Red Drum calling to peak after sunset [17, 23], it is conceivable that fish may use other times to call when vessel noise is not as intense. When recording aquacultured Red Drum, researchers noted fish calling peaked in the morning hours [53], though this was not attributed to disturbance from noise pollution. Though this is outside the context of our study, assessing the full 24h cycle of sound production in the face of vessel noise may be worth researching in the future.

This research also adds to the ever-growing literature that shows anthropogenic noise is highly prevalent in marine coastal habitats [3, 9, 54]. We documented significant periods of noise pollution in the environment with fish sounds over repeated spawning seasons (Fig. 7). Noise pollution has been seen in other aquatic systems and is concerning in its prevalence. In May River, South Carolina researchers saw a 21% overlap in time for Red Drum calls and vessel noise [28]. Notably in Oct. 2021, our study showed on average 100% of the CP contained either vessel noise or Red Drum calling, which means every opportunity for fish to call without potential masking of their calls was used. This leads to the reasonable conclusion that significant overlap in time exists between vessel noise and when Red Drum calling occurs in Saint Andrew Bay. Beyond the potential for altered acoustic behavior, this excessive noise exposure could present further risk to Red Drum and other marine life.

## Conclusion

Anthropogenic noise poses a serious risk to animals’ ability to perform basic tasks by distorting the natural soundscape. Foraging behavior, predator detection and avoidance, and interaction with conspecifics can all be altered due to noise pollution [55]. Moreover, noise is a physiological stressor [2] and is well established as a cause of hearing loss for some aquatic animals [4, 56]. The ability to perceive sound and distinguish between relevant acoustic signals in the environment is directly tied to an individual’s fitness [57]. Additionally, coastal habitats are biologically productive and used by many species at different life stages for foraging, shelter, migration, spawning, and nursery habitat [58]. Though, while critically important for these species, nearshore habitats experience disproportionate anthropogenic impacts as human activity is coastally concentrated [5–7, 51, 59]. In addition, the ability of sound to impact large regions from its source makes it ever more critical to assess implications of vessel noise in coastal waters, as its effects may be widespread and varied, even in similar taxa occupying shared habitat [40, 47]. Continued investigation of sciaenid vessel noise impacts is warranted, with both lab experimentation and in situ studies. Assessment of sound production by individual fish or small groups in controlled environments will allow for greater insights into how these animals respond to noise pollution. Ongoing use of passive acoustic monitoring in coastal regions is needed to elucidate multispecies responses to the ever-growing noise pollution found in these regions. A comprehensive understanding of noise pollution impacts on aquatic life is critical for coastal ecosystem management.

## Acknowledgements

We thank David Muncher (Louisiana Universities Marine Consortium) and Neal Kolonay for supporting deployments by scuba and assistance in the field. We also thank Bryce Newman for helping with assistance on data analysis. For access to recording sites, we thank the Florida Department of Environmental Protection (permit # 02252022011), Saint Andrews State Park.

## Author contribution

Conceived and designed the experiments: BHP KSB. Performed the experiments: BHP AK TEC KSB. Analyzed the data: BHP DB KSB. Contributed reagents/materials/analysis tools: KSB TEC. Wrote the paper: BHP KSB.

## References

1. McCauley RD, Fewtrell J, Popper AN. High intensity anthropogenic sound damages fish ears. J Acoust Soc Am. 2003;113(1):638–42.

2. Smith ME, Kane AS, Popper AN. Noise-induced stress response and hearing loss in goldfish (Carassius auratus). J Exp Biol. 2004;207(Pt 3):427–35.

3. Duarte CM, Chapuis L, Collin SP, Costa DP, Devassy RP, Eguiluz VM, et al. The soundscape of the Anthropocene ocean. Science. 2021;371(6529):eaba4658.

4. Badlowski GA, Boyle KS. Repeated boat noise exposure damages inner ear sensory hair cells and decreases hearing sensitivity in Atlantic croaker (Micropogonias undulatus). J Exp Biol. 2024;227(2):jeb245093.

5. Picciulin M, Sebastianutto L, Codarin A, Calcagno G, Ferrero EA. Brown meagre vocalization rate increases during repetitive boat noise exposures: a possible case of vocal compensation. J Acoust Soc Am. 2012;132(5):3118–24.

6. La Manna G, Manghi M, Perretti F, Sarà G. Behavioral response of brown meagre (Sciaena umbra) to boat noise. Mar Pollut Bull. 2016;110(1):324–34.

7. de Jong K, Amorim MCP, Fonseca PJ, Fox CJ, Heubel KU. Noise can affect acoustic communication and subsequent spawning success in fish. Environ Pollut. 2018;237:814–23.

8. Vieira M, Amorim MC, Fonseca PJ. Fish sound production and vocalization behavior. In: Slabbekoorn H, editor. Effects of anthropogenic noise on animals. New York: Springer; 2021.

9. Hildebrand J. Anthropogenic and natural sources of ambient noise in the ocean. Mar Ecol Prog Ser. 2009;395:5–20.

10. Wilson L, Pine MK, Radford CR. Small recreational boat: a ubiquitous source of sound pollution in shallow coastal habitats. Mar Pollut Bull. 2022;174:113295.

11. Rogers PH, Cox M. Underwater sound as a biological stimulus. In: Atema J, Fay RR, Popper AN, editors. Sensory biology of aquatic animals. New York: Springer; 1988. p. 131–49.

12. Amoser S, Wysocki LE, Ladich F. Noise emission during the first powerboat race in an Alpine lake and potential impact on fish communities. J Acoust Soc Am. 2004;116(6):3789–97.

13. Ramcharitar J, Gannon DP, Popper AN. Bioacoustics of fishes of the family Sciaenidae (croakers and drums). Trans Am Fish Soc. 2006;135(5):1409–31.

14. Ladich F. Ecology of sound communication in fishes. Fish Fish. 2019;20(3):552–63.

15. Connaughton MA, Taylor MH, Fine ML. Effects of fish size and temperature on Weakfish disturbance calls: implications for the mechanism of sound generation. J Exp Biol. 2000;203(Pt 10):1503–12.

16. Gannon DP. Acoustic behavior of Atlantic Croaker, Micropogonias undulatus (Sciaenidae). Copeia. 2007;2007(1):193–204.

17. Lowerre-Barbieri SK, Barbieri LR, Flanders JR, Woodward AG, Cotton CF, Knowlton MK. Use of passive acoustics to determine Red Drum spawning in Georgia waters. Trans Am Fish Soc. 2008;137(2):562–75.

18. Barros NB, Wells RS. Prey and feeding patterns of resident bottlenose dolphins (Tursiops truncatus) in Sarasota Bay, Florida. J Mammal. 1998;79(3):1045–59.

19. Chao NL, Frédou FL, Haimovici M, Peres MB, Polidoro B, Raseira M, et al. A popular and potentially sustainable fishery resource under pressure—extinction risk and conservation of Brazilian Sciaenidae. Glob Ecol Conserv. 2015;4:117–26.

20. Anderson J, McDonald D, Bumguardner B, Olsen Z, Ferguson JW. Patterns of maturity, seasonal migration, and spawning of Atlantic Croaker in the western Gulf of Mexico. Gulf Mex Sci. 2018;34(1–2):19–31.

21. National Marine Fisheries Service. Fisheries of the United States, 2020. Silver Spring (MD): NOAA; 2022. Current Fishery Statistics No. 2020.

22. Montie EW, Kehrer C, Yost J, et al. Long-term monitoring of captive red drum Sciaenops ocellatus reveals that calling incidence and structure correlate with egg deposition. J Fish Biol. 2016;88:1776–95.

23. Monczak A, McKinney B, Souiedan J, et al. Sciaenid courtship sounds correlate with juvenile appearance and abundance in the May River, South Carolina, USA. Mar Ecol Prog Ser. 2022;693:1–17.

24. Hightower CL, Drymon JM, Powers SP. Current Status of Adult Red Drum (Sciaenops ocellatus) in the North Central Gulf of Mexico. SEDAR49-DW-16. North Charleston (SC): SEDAR; 2016.

25. Fish M, Mowbray W. Sounds of Western North Atlantic fishes. Baltimore: The Johns Hopkins Press; 1970.

26. Holt SA. Distribution of Red Drum spawning sites identified by a towed hydrophone array. Trans Am Fish Soc. 2008;137(2):551–61.

27. Horodysky AZ, Musick JA, Latour RJ. Baseline thresholds of hearing in sciaenid fishes. J Acoust Soc Am. 2008;123(5):3832.

28. Smott S, Monczak A, Miller ME, Montie EW. Vessel noise impacts on sciaenid fishes of the northern Gulf of Mexico. J Acoust Soc Am. 2018;144(3):1749.

29. Florida Fish and Wildlife Conservation Commission. 2024 Panhandle Redfish Annual Review. Tallahassee: FWC; 2024.

30. Lombard E. Le signe de l’élévation de la voix [The sign of the elevation of the voice]. Ann Malad Oreille Larynx. 1911;37:101–9.

31. United States Naval Observatory. Astronomical Applications Department [Internet]. Washington (DC): USNO; 2024. Available from: USNO Portal.

32. K. Lisa Yang Center for Conservation Bioacoustics. Raven Lite: Interactive Sound Analysis Software [software]. Version 2.0.5. Ithaca (NY): The Cornell Lab of Ornithology; 2024. Available from: Raven Software.

33. R Core Team. R: A language and environment for statistical computing [software]. Version 4.4.1. Vienna, Austria: R Foundation for Statistical Computing; 2024. Available from: R Project.

34. Sueur J, Aubin T, Simonis C. Seewave, a free modular tool for sound analysis and synthesis. Bioacoustics. 2008;18(2):213–26.

35. Ma BB, Nystuen JA, Lien RC. Prediction of underwater sound levels from rain and wind. J Acoust Soc Am. 2005;117(6):3555–65.

36. Pinheiro J, Bates D, R Core Team. nlme: Linear and Nonlinear Mixed Effects Models. R package version 3.1–164; 2023. Available on CRAN.

37. Burnham KP, Anderson DR. Model Selection and Multimodel Inference: A Practical Information-Theoretic Approach. 2nd ed. New York: Springer; 2002.

38. Pinheiro JC, Bates DM. Mixed-Effects Models in S and S-PLUS. New York: Springer-Verlag; 2000.

39. Zollinger SA, Brumm H. The Lombard effect. Curr Biol. 2011;21(16):R614–5.

40. Matos F, Amorim MC, Fonseca PJ, Vieira M. Reaction of two sciaenid species to passing boats. Estuar Coast Shelf Sci. 2024;299:108643.

41. Luczkovich JJ, Krahforst CS, Sprague MW. Does Vessel Noise Change the Calling Rate and Intensity of Soniferous Fishes? In: Popper A, Hawkins A, editors. The Effects of Noise on Aquatic Life. New York: Springer; 2012. p. 375–8.

42. Picciulin M, Zucchetta M, Facca C, Malavasi S. Boat-induced pressure does not influence breeding site selection of a vulnerable fish species. Mar Pollut Bull. 2022;180:113750.

43. Luczkovich JJ, Krahforst CS, Kelly KE, Sprague MW. The Lombard effect in fishes: How boat noise impacts oyster toadfish vocalization amplitudes. POMASA. 2016;27:010035.

44. Brown NAW, Halliday WD, Balshine S, Juanes F. Low-amplitude noise elicits the Lombard effect in plainfin midshipman mating vocalizations in the wild. Anim Behav. 2021;181:29–39.

45. Ceraulo M, Sal Moyano MP, Hidalgo FJ, et al. Boat Noise and Black Drum Vocalizations in Mar Chiquita Coastal Lagoon (Argentina). J Mar Sci Eng. 2021;9(1):44.

46. Price BP, Boyle KS. Disruption of Red Drum calling by vessel noise. Mar Environ Res. 2025;211:107449. [Note: In bibliography as Boyle et al. 2025].

47. Boyle KS, Cox TE, Kirkland AM, Price BP. Influences and interactions of vessel noise and environmental variables on Silver Perch and seatrout spawning choruses. Mar Pollut Bull. 2025;219:118228.

48. Pierce AD. Acoustics: An Introduction to its Physical Principles and Applications. New York (NY): Acoustical Society of America; 1989.

49. Sprague MW, Luczkovich JJ. Measurement of an individual silver perch Bairdiella chrysoura sound pressure level in a field recording. J Acoust Soc Am. 2004;116(5):3186–91.

50. Parsons JG, McCauley RD, Mackie MC, et al. In situ levels of mulloway (Argyrosomus japonicus) calls. J Acoust Soc Am. 2012;132(5):3559–68.

51. Picciulin M, Sebastianutto L, Codarin A, Calcagno G, Ferrero EA. Brown meagre vocalization rate increases during repetitive boat noise exposures. J Acoust Soc Am. 2012;132(5):3118–24.

52. Somogyi NA, Rountree RA. The sound production of Aplodinotus grunniens in the presence of boat sounds. J Acoust Soc Am. 2023;154(2):831–40.

53. Parmentier E, Tock J, Falguière JC, Beauchaud M. Sound production in Sciaenops ocellatus: Preliminary study for the development of acoustic cues in aquaculture. Aquaculture. 2014;432:204–11.

54. Slabbekoorn H, Bouton N, van Opzeeland I, et al. A noisy spring: the impact of globally rising underwater sound levels on fish. Trends Ecol Evol. 2010;25(7):419–27.

55. Simpson SD, Radford AN, Nedelec SL, et al. Anthropogenic noise increases fish mortality by predation. Nat Commun. 2016;7:10544.

56. Smith ME, Schuck JB, Gilley RR, Rogers BD. Structural and functional effects of acoustic exposure in goldfish: evidence for tonotopy in the teleost saccule. BMC Neurosci. 2011;12:19.

57. Fay RR, Popper AN. Evolution of hearing in vertebrates: The inner ears and processing. Hear Res. 2000;149(1–2):1–10.

58. Seitz RD, Wennhage H, Bergstrom U, et al. Ecological value of coastal habitats for commercially and economically important species. ICES J Mar Sci. 2014;71(3):648–65.

59. Crossett K, Ache B, Pacheco P, Haber K. National Coastal Population Report, Population Trends from 1970 to 2020. Washington (DC): NOAA; 2013.

60. Holles S, Simpson S, Radford A, et al. Boat noise disrupts orientation behaviour in a coral reef fish. Mar Ecol Prog Ser. 2013;485:295–300.

61. Mok HK, Wu SC, Sirisuary S, Fine ML. A sciaenid swim bladder with long skinny fingers produces sound with an unusual frequency spectrum. Sci Rep. 2020;10:18619.

62. Smith ME, Monroe JD. Causes and consequences of sensory hair cell damage and recovery in fishes. In: Sisneros JA, editor. Fish hearing and bioacoustics. Switzerland: Springer; 2016. p. 393–417.

63. Tower R. The production of sound in the drumfishes, the searobin, and the toadfish. Ann N Y Acad Sci. 1908;18(1):149–80.

64. Vance TL, Hewson JM, Modla S, Connaughton MA. Variability in sonic muscle size and innervation among three sciaenids: spot, Atlantic croaker, and weakfish. Copeia. 2002;(4):1137–43.

